# Migration of gastric cancer is suppressed by rabies virus glycoprotein via regulating α7-nicotinic acetylcholine receptors/ERK- EMT

**DOI:** 10.1101/497362

**Authors:** Xufeng Bu, Anwei Zhang, Zhengwei Chen, Xunfeng Zhang, Chao yun Yin, Jie Zhang, Yao Zhang, Riting Zhang, Xiaomei Shao, Yulan Yan

## Abstract

Nicotinic acetylcholine receptors (nAChRs) have been reported to be overexpressed in malignancies in humans and is associated with tumorigenesis and cell migration. In previous studies of gastric cancer, alpha7 nicotinic acetylcholine receptor (α7-nAChR) overexpression can induce epithelial-mesenchymal transition (EMT) and promote migration of gastric cancer cells. Recombinant avirulent Newcastle disease virus (NDV) LaSota strain expressing the rabies virus glycoprotein (rL-RVG) may promote apoptosis of gastric cancer cells and reduces migration of lung cancer metastasis. However, whether rL-RVG inhibits migration of gastric cancer cells and what the underlying functional mechanism is remains unknown. In this study, our findings demonstrate that rL-RVG suppressed migration and reduced EMT of gastric cancer cells via α7-nAChR in vitro. Furthermore rL-RVG decreased the phosphorylation levels of the MEK/ERK signaling pathway such as down-regulating the expression of P-MEK and P-ERK. Additionally, rL-RVG also reduced the expression level of mesenchymal markers N-cadherin and Vimentin and enhanced the expression of the epithelial marker E-cadherin. Lastly, rL-RVG together with nicotinic acetylcholine receptors (nAChRs) inhibited gastric cancer epithelial to mesenchymal transition (EMT) which suppressed gastric cancer cell migration. We also found that rL-RVG suppresses the growth of gastric cancer subcutaneous tumor cells *in vivo*. Thus, rL-RVG inhibits α7-nAChR-MEK/ERK-EMT to suppress migration of gastric cancer cells.

## Introduction

Stomach cancer is the fifth most common malignancy and the third major cause in cancer-related deaths in the word ^[1]^. Moreover, the death rate of stomach cancer is the second highest among all cancer related deaths in China ^[2]^. It is difficult to detect the early stages of gastric cancer and there are an overwhelming number of patients who are diagnosed at the advanced stages of disease ^[3]^. Therefore, the study of the mechanisms of cancer cell migration mechanism during stomach cancer is of great significance for new drug discoveries and for the development of effective treatments for gastric cancer. Currently, the main treatment of gastric cancer consists only of surgical treatment, radiotherapy and chemotherapy. However, these methods are accompanied by poor patient prognosis with advanced gastric cancer and therefore a novel treatment is urgently needed for people affected by this devastating disease.

Nicotinic acetylcholine receptors (nAChRs) are a type of ligand-gated ion channel proteins, and their expression is not only found in neuronal cells but also in non-neuronal cells including gastric cancer cells ^[4,5]^. Alpha7 nicotinic acetylcholine receptor (α7-nAChR) is a member of the family of nAChRs and is widely distributed in epithelial cells of the stomach ^[6,7]^. It has been reported that the activation of α7-nAChR plays an important role in the proliferation and migration of cancer cells. Extracellular signal-regulated kinase (ERK) signaling pathway is involved in a variety of functions and activated by several stimuli including α7-nAChR^[8]^.

Epithelial to mesenchymal transition (EMT) is the original biological step required for the invasion of cancer cells and metastasis. A hallmark of EMT includes the presence of mesenchymal markers such as N-cadherin, Vimentin and epithelial marker E-cadherin^[9,10]^. EMT is of great significance in tumor migration ^[11]^, and a recent study also suggested that alpha 7-nAChR overexpression could enhance EMT to promote proliferation and migration of gastric cancer cells^[12]^. Moreover, it was reported that nicotine could promote EMT to induce migration of cancer cells via regulation of the alpha 7-nAChR/MEK/ERK pathway ^[13]^.

Alpha 7-nAChR may be a potential therapeutic key point for the treatment of stomach cancer. Previous studies have also demonstrated that recombinant avirulent NDV LaSota strain, expressing the rabies virus glycoprotein (rL-RVG), could induce the apoptosis of gastric cancer cells as well as suppress the migration of lung cancer cells by regulating α7-nAChR ^[14,15]^. However, it still remains unclear whether rL-RVG could also suppress the migration of gastric cancer and its underlying mechanism. In this study, we would explored if rL-RVG could suppress migration of gastric cancer via regulating α7-nAChR/ERK signaling and EMT.

## Materials and methods

### Materials

The rL-RVG, NDV, anti-RVG antibody and anti-NDV antibody were stored at −80°C supplied by the Harbin Veterinary Research Institute (Harbin, China). The gastric cancer cell lines SGC7901 and BGC were purchased from biological sciences cell resource center of the Chinese academy of sciences, Shanghai Institute. Nude mice were purchased from the Animal Experiment Center of Yangzhou University (Yangzhou, China). Methyllcaconitine citrate hydrate (MLA) (5mg/mL) which worked as a specific competitive α7-nAChR antagonist was purchased from Santa Cruz (California, USA). Acetylcholine bromide (ACB) (5mg/mL), an acetylcholine agonist, was obtained from Sigma-Aldrich (St. Louis, MO, USA). In addition, small interfering RNA (si-RNA) for α7-nAChR was purchased from RiboBio (GuangZhou, China). Corynoxenine (10mM 1mL in DMSO), the inhibitor of the MEK-ERK pathway, was purchased from MedChemExpress (MCE, USA). Rabbit polyclonal anti-α7 nAChR was purchased from Abcam (London, UK); rabbit polyclonal anti-MEK1/2, anti-P-MEK1/2, anti-ERK1/2, anti-P-ERK1/2 and anti-snail were purchased from Cell Signaling Technology (CST, USA); Mouse monoclonal anti-N-Cadherin, E-Cadherin and Vementin were purchased from Boster (WuHan, China).

### Cell Culture Reagents, Viruses and Treatment

The cell lines BGC and SGC7901 were cultured in RPMI 1640 supplemented with 10% fetal bovine serum, penicillin (100U/mL), and streptomycin (100μg/mL) in a humidified incubator (37°C, 5%CO_2_). In addition, when BGC and SGC cell lines reached their logarithmic proliferation stage of up to 80% confluency, they were then sub cultured or used for experiments. The cultured cell lines were randomly divided into rL-RVG, NDV and PBS groups, along with the MLA, ACB, si-RNA of α7-nAChR and corynoxenine pretreatment groups.

### CCK-8 Assay

The viability of infected BGC and SGC were monitored by performing a CCK-8 assay. BGC and SGC were harvested, pelleted by centrifugation and counted using a blood counting chamber. Cells (6×10^3^) were then seeded into 96-well plates and grown in media containing varying dilution titers of rL-RVG or NDV 24 h. The CCK-8 reagent was added into each well incubated for another 4 h. Lastly, the color of the media changed in each well and was measured at 450 nm using a spectrophotometer.

### Clonogenic survival assay

BGC and SGC were seeded into 6-well plates (1000 cells per well) and then infected with rL-RVG, NDV at a multiplicity of infection of 10 or with PBS for 24 h. After incubation in media for 10 days, the cells were fixed by absolute ethyl alcohol for 30 minutes and stained for 1 hour with crystal violet (0.2%) to visualize cell colonies. Each individual experiment was repeated three times.

### Wound-Healing Assay

BGC and SGC were added into 6-well plates and randomly divided into rL-RVG, NDV and PBS treated groups in which cell were infected by rL-RVG, NDV or PBS for 24 h. Groups pretreated with ACB, MLA, si-RNA or corynoxeine for 24 h were also set up. When cells reached 80% confluency, cell monolayers were wounded with a 10 μL pipette tip after 24 h and their wound healing ability was observed by microscopy.

### Migration Assay

To carry out migration assay 24-well plate transwell units with polycarbonate membrane (8.0μm pore size; Costar, MA, USA) were used for in-vitro experiments according to the manufacturer’s protocol. In short, the upper chamber of filter inserts contained serum free medium with 1×10^5^ cells belonging to one of the treatment groups: rL-RVG, NDV, PBS, MLA, ACB, si-RNA of α7-nAChR or corynoxenine. Meanwhile, the lower chamber was filled with 600 μL 10% serum media. After incubation at 37°C in 5%CO_2_ for 24 h, the media in the lower chamber were collected and then non migrating cells attached to the membrane of the upper chamber surface were scrubbed. Finally, cells on the bottom wells were stained with 0.1% crystal violet for 20 minutes, and then washed with PBS for 3 times. Lastly, the migrating cells were counted under the microscope and analyzed by using Image-J software. (National Institutes of Health, USA).

### Western Blot Analysis

After pre-treatment with MLA or ACB or si-RNA of α7-nAChR or corynoxenine for 12 h，cells were infected for 24 h with either rL-RVG, NDV or PBS and then washed with ice-cold PBS for 3 times and lysed by using the lysis buffer RIPA containing 1 mM PMSF for 30 minutes on ice. Next the lysates were collected and the protein concentrations were quantified using a BCA kit (Thermo Fisher Scientific, USA). Equal quantities of protein were separated by using a 10% SDS-PAGE and the proteins were then transferred to polyvinylidinedifluoride (PVDF) membranes (Bio-Rad Laboratories). The membranes were then blocked with 5% BSA in Tris-buffered saline containing 0.1% Tween 20 (TBST, at pH 7.5) for about 2 hours at room temperature before washing them with TBST for 15 minutes for 3 times. Next the membranes were incubated with antibodies at 4°C overnight with the following antibodies: anti-α7 nAChR, anti-P-MEK, anti-MEK, anti-P-ERK, anti-ERK, anti-E-cadherin, anti-N-cadherin and anti-Vementin. Proteins were detected with HRP-conjugated secondary antibodies for 1 h at room temperature. The protein bands were visualized with a Typhoon 9400 variable mode imager (Amersham Biosciences, UK) using chemiluminescence (ECL Plus Substrate, Thermo Fisher Scientific, USA).

### Immunofluorescent Assay

BGC and SGC cells were added into 24-well plates and fixed with 4% paraformaldehyde for 2 h at room temperature. Cells were then permeabilized with 0.5 % TritonX-100 in PBS for 10 minutes and blocked with 5% BSA for 1 h. Next the cells were washed 3 times with PBS for 5 min before incubating them with anti-P-ERK at 4°C overnight. On the next day, the secondary antibody was used to stain cells in order to be detected for immunofluorescence microscopy.

### Xenografts

After infection of BGC, SGC cells with rL-RVG, NDV and PBS for 24 h, the cells were subcutaneously injected into axillary subcutaneous tissues of adult female athymic-nude mice which were randomly selected for the treatment with either rL-RVG, NDV or PBS and housed under specific pathogen free conditions. The size of subcutaneous tumors that developed were measured 2 weeks post treatment using the following calculation: V=W^2^L0.5 (V is volume, W is width and L is length).

### Statistical Analysis

All collected experimental data is presented as mean ± SD. A Student’s t-test or one-way ANOVA with Bonferroni post-test was used to calculate statistical significance using the GraphPad Prism7.0 software (La Jolla, CA, USA). A p-value of *P<0.05* or *P<0.01* was considered statistically significant. Each experiment was conducted independently and repeated at least 3 times.

## Results

### RVG and NDV protein expression in gastric cancer cells

To investigate the mechanism of rL-RVG suppressing the migration of gastric cancer cells, we first analyzed the expression of rL-RVG and NDV proteins in gastric cancer. Previous studies show that lung cancer cell exhibit a stable expression of RVG and NDV proteins by PCR, Western blot and immunofluorescence microscopy ^[16]^. In our study, we used Western blot to analyze both RVG and NDV protein expression in virally infected gastric cancer cells and found that RVG proteins were only expressed in the rL-RVG group while the expression of NDV proteins was expressed in both the rL-RVG and NDV group (Fig. 1A).

**Figrure 1.**
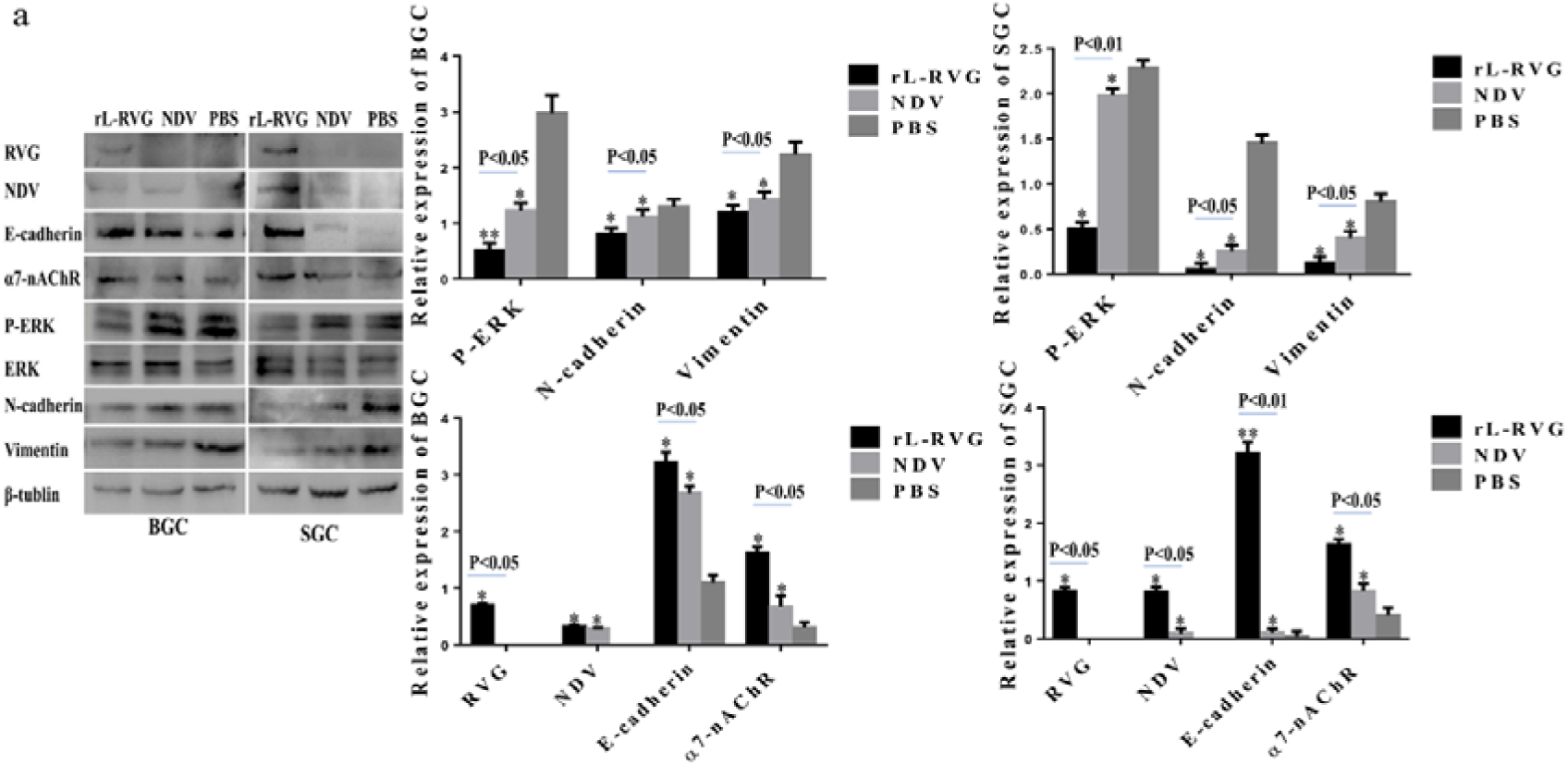
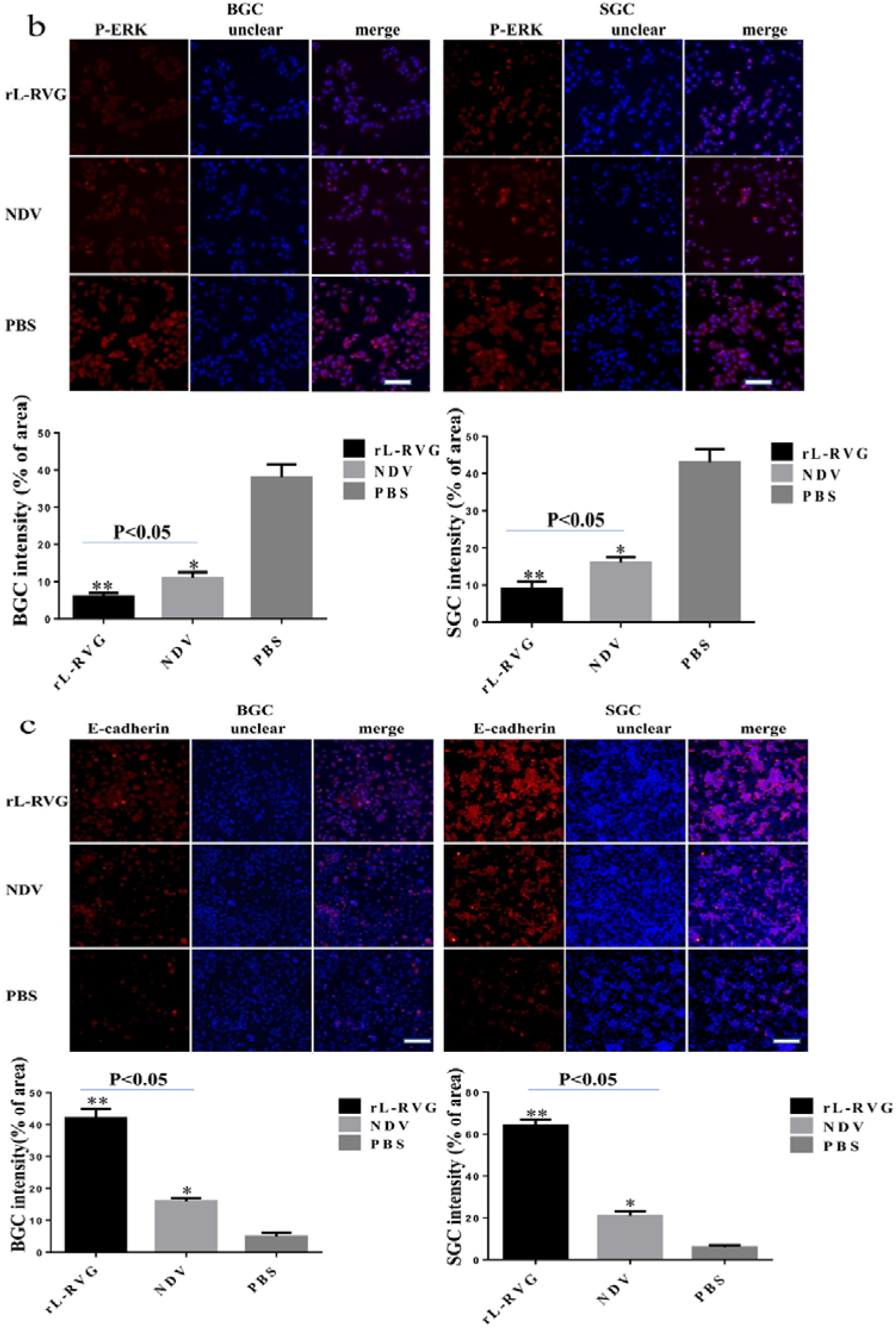
Expression of RVG, NDV, α7-nAChR, MEK/ERK signaling pathway and epithelial/mesenchymal markers proteins in infected BGC and SGC cells. **a**. Western blot analysis of RVG, NDV, α7-nAChR, MEK/ERK signaling pathway and epithelial/mesenchymal proteins. **b**. Immunofluorescence analysis of P-ERK. **c**. Immunofluorescence analysis of EMT protein markers E-cadherin. BGC and SGC cells were infected with either rL-RVG, NDV and PBS for 24 h. *\*P<0.5*, *\*\*P<0.01*.(rL-RVG vs NDV and PBS groups, respectively, Bar=25 μm).

### rL-RVG suppressed the proliferation and migration of gastric cancer cells

Metastasis is a big hurdle in the management of gastric cancer. To study metastasis migration we used a transwell based and wound healing assay to monitor the influence of viruses on gastric cancer cells migration. After infecting cells with rL-RVG or NDV for 24 h, we observed that rL-RVG and NDV both reduced the migration of gastric cancer cells compared to the PBS treated group. Of note is that the inhibitory migration was stronger in rL-RVG treated cells compared to the NDV group (Fig 2A,B). Moreover, we found that rL-RVG had an inhibitory influence on the migration of both SGC and BGC and SGC-7901 cells and were therefore selected for further analysis in subsequent experiments.

**Figure 2.**
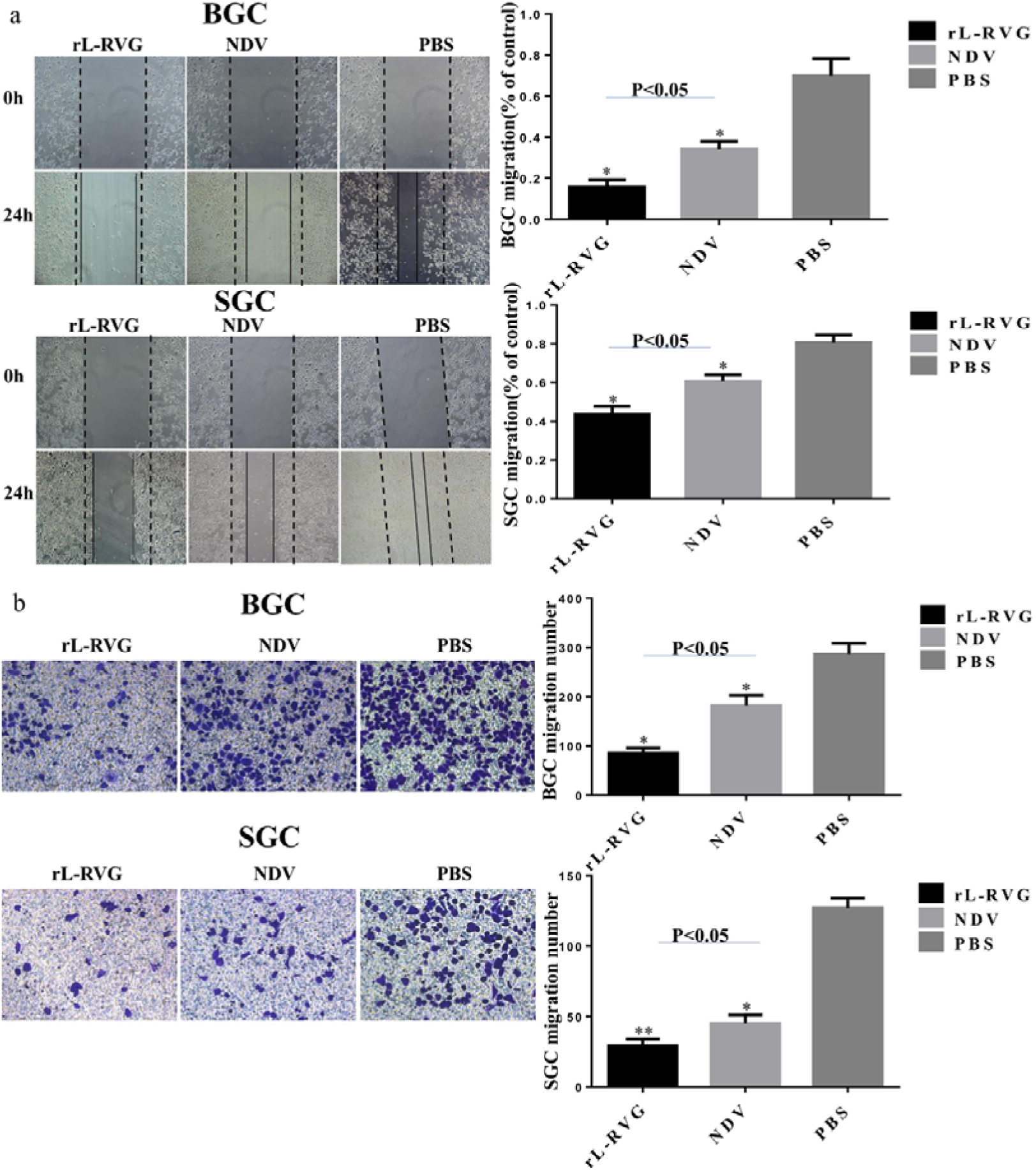
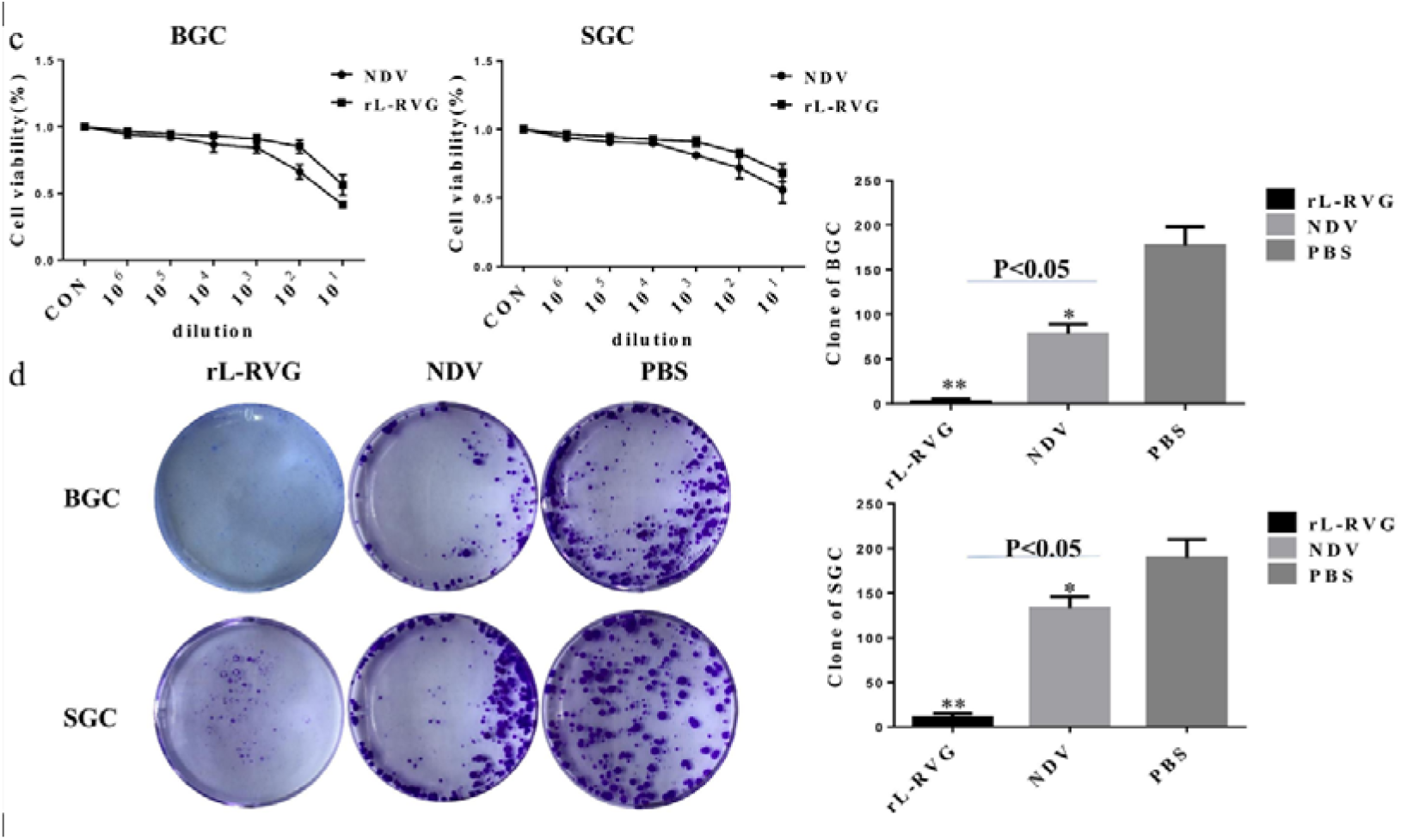
rL-RVG suppresses the proliferation and migration of BGC and SGC cells. **a**. Healing and **b**. transwell assays were used to monitore the migration of BGC and SGC cells infected with rL-RVG, NDV and PBS, respectively. **c**. Influence of different rL-RVG, NDV dilution titers on the viability of BGC and SGC cells. **d**. The clonogenic activity of BGC and SGC cells after infection with rLRVG and NDV at a multiplicity of infection of 10. Colony formation was attenuated in the rL-RVG group. *\*P<0.5*, *\*\*P<0.01*.(rL-RVG vs NDV and PBS groups, respectively).

To detect the viability of gastric cancer cells SGC7901 and BGC cells were infecting with rL-RVG or NDV for 24 h and analysed using a CCK8 assay. rL-RVG and NDV both suppressed cell proliferation in a concentration-dependent manner but overall rL-RVG had a stronger inhibitory effect on proliferation compared to NDV and the PBS control group. If rL-RVG and NDV were diluted to 10_3_ and 10_2_ respectively, the viability of SGC and BGC cells was higher than 80%, and the viral titer was approximately 10^9.8^ EID_50_ /mL (Fig 2C,D).

### The role of α7-nAChR in the process of rL-RVG regulated migratory abilities of gastric cancer cells

The expression of α7-nAChR in the rL-RVG and NDV groups was higher compared with PBS group using Western Blot analysis. However the cell migratory ability in both the rL-RVG and NDV group was suppressed compared with the PBS control blank group. rL-RVG suppressed the migration more potently compared with the NDV and PBS groups (Fig 1A, Fig 2A,B). Further exploration regarding the role of α7-nAChR in rL-RVG on the suppression of the migration of gastric cancer cells is necessary and the following treatments were performed.

MLA, an antagonist of α7-nAChR was used to pre-treat SGC7901 cells for 24 h before infecting them with virus. Our result show that α7-nAChR expression was inhibited in a competitive manner and moreover, the migration of SGC7901 cells was more suppressed in the MLA pre-treatment groups compared to the not pre-treated groups as shown in our migration and wound healing assays. This result suggests that rL-RVG may suppress cell migration through the α7-nAChR pathway (Fig 4A,B).

To further verify the role of α7-nAChR in rL-RVG-induced cells migration, we used small interfering RNA methods to knock down the expression of α7-nAChR. We found that our results were consistent with the groups pre-treated with MLA. Therefore, rL-RVG may play a role as competitive antagonist of α7-nAChR to inhibit the migration of SGC7901 cells (Fig 5A,B).

In support of these results we found that ACB, an agonist of α7-nAChR stimulates the expression of α7-nAChR (Fig 3A,B). In contrast to this, we got the opposite result when comparing our result with MLA or si-RNA pre-treated cells. Thus, rL-RVG suppresses the migration of SGC7901 cells by inhibiting α7-nAChR competitively.

**Figure 3.**
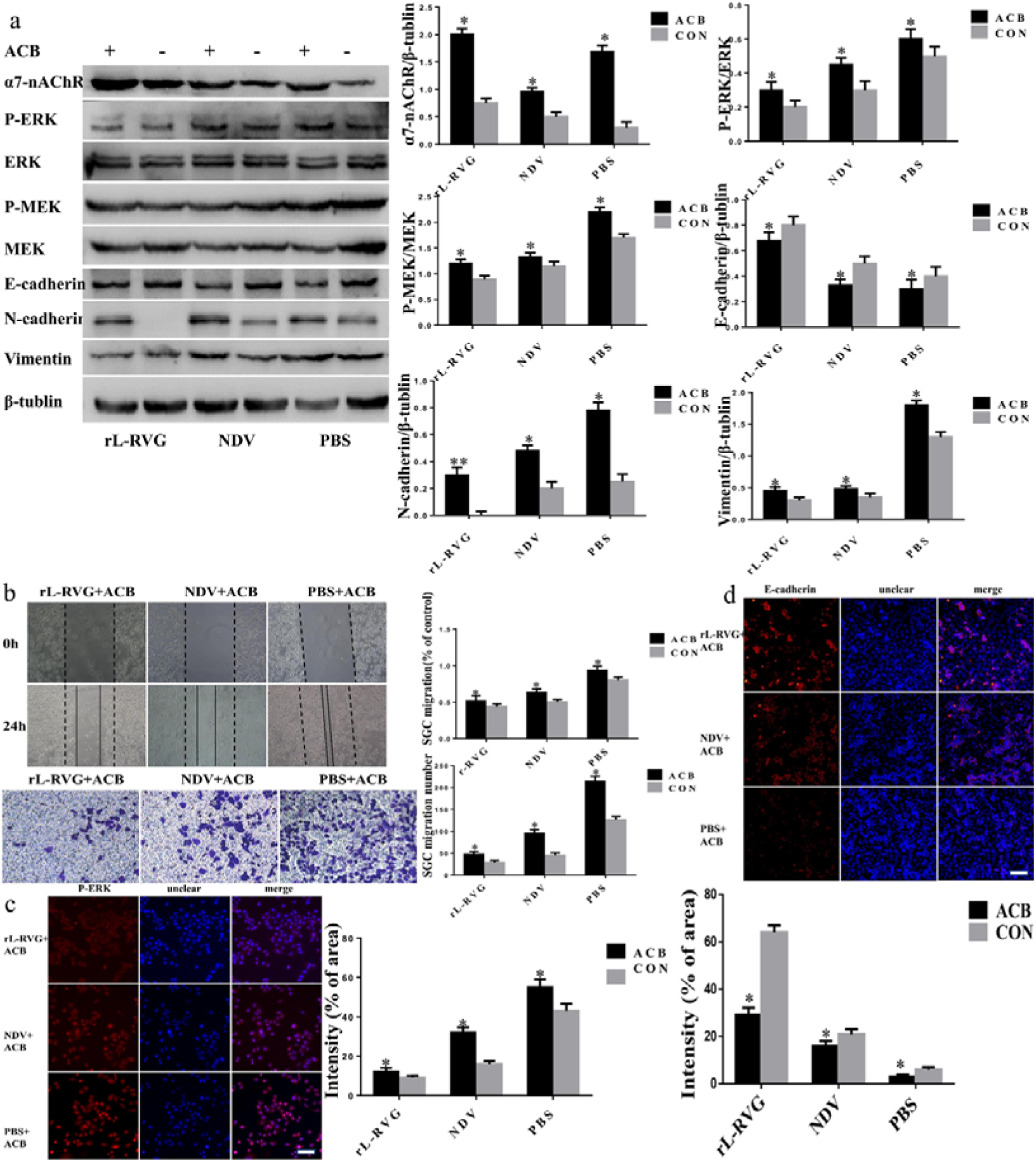
Effects of rL-RVG and ACB pretreated SGC cells on the α7-nAChR, MEK/ERK signaling pathway, epithelial/mesenchymal proteins and cell migration. **a**. Western blot analysis of α7-nAChR, MEK/ERK signaling pathway and epithelial/mesenchymal protein marker. **b**. Cell migration was detected by wound healing and transwell assay. **c**. Immunofluorescence analysis of P-ERK. **d**. Immunofluorescence analysis of EMT proteins E-cadherin in infected SGC cells. *\*P<0.5, **P<0.01*.(rL-RVG+ACB vs rL-RVG, NDV+ACB vs NDV, PBS+ACB vs PBS, respectively, Bar=25 μm).

### MEK/ERK signaling pathway was involved in rL-RVG-lowing migration of gastric cancer cell

It was shown previously that α7-nAChR activates several signaling pathways in connection with tumorigenic effects including the MEK/ERK pathway^[8]^. The role of the extracellular signal-regulated kinase (ERK) signaling pathway in the rL-RVG-induced suppression of cell the migration during gastric cancer remains unclear. Western blot analysis showed that rL-RVG induces the down-regulation of phosphorylation levels of ERK1/2 when α7-nAChR expression was blocked by rL-RVG. Furthermore the migratory ability of gastric cancer cells was also reduced. Consistent with these results we showed by immunofluorescence that the expression level of P-ERK was down-regulated after being infected with either rL-RVG or NDV but was lower in the rL-RVG group compared with the NDV and PBS blank control group (Fig 1 A,B and Fig 2 A,B).

To further support our results we showed that the level of ERK1/2 phosphorylation was lower compared with the non pretreated groups after prtreatment with MLA or si-RNA of α7-nAChR by both, Western blot and immunofluorescent assay (Fig 4A,C and Fig 5A,C). Our migration and wound healing assay showed that the migratory ability of SGC cells in the pretreated group decreased more compared to non pretreated cells (Fig 4B and Fig 5B). To clarify whether rL-RVG modulates the MEK/ERK signaling pathway to suppress cell migration, corynoxenine, an inhibitor of the MEK/ERK pathway, was used to pre-treat gastric cancer cells (Fig 6A-C). We obtained comparable results in the groups treated with MLA and si-RNA of α7-nAChR as described above. However, opposite results were obtained when cells were pre-treated with ACB (Fig 3A-C). These results indicate that rL-RVG attenuates the activation levels of MEK/ERK pathway via blocking α7-nAChR by competition.

**Figure 4.**
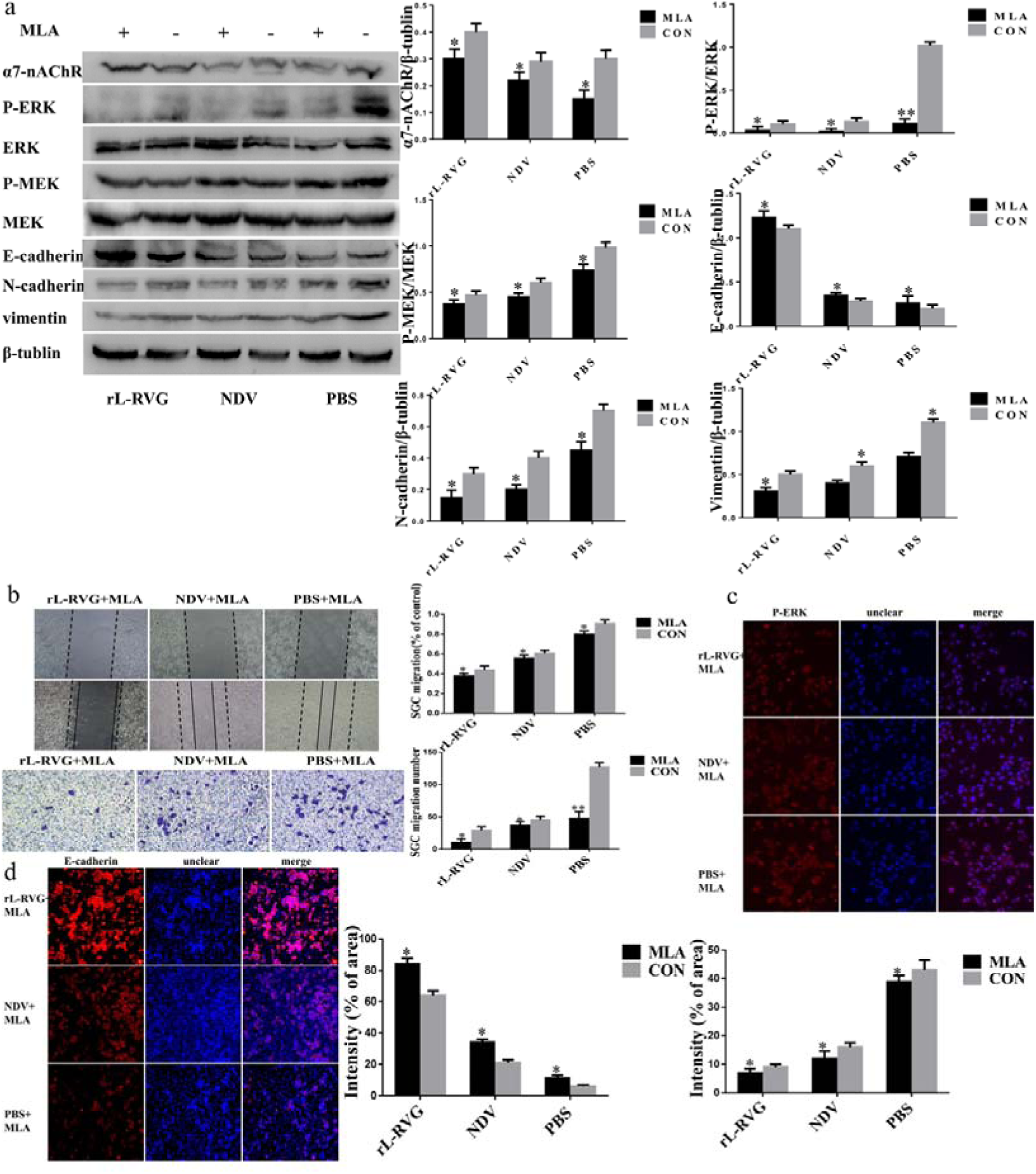
Effects of rL-RVG and MLA pretreated cells on α7-nAChR, MEK/ERK signaling pathway and epithelial/mesenchymal proteins and migration of cells. **a**. Western blot analysis of the α7-nAChR, MEK/ERK signaling pathway and epithelial/mesenchymal protein markers. **b**. Cell migration was detected by wound healing and transwell assay. **c**. Immunofluorescence analysis of P-ERK. **d**. Immunofluorescence analysis of EMT markers E-cadherin in infected SGC cells. *\*P<0.5, **P<0.01* (rL-RVG+MLA vs rL-RVG, NDV+MLA vs NDV, PBS+MLA vs PBS, Bar=25 μm).

**Figure 5.**
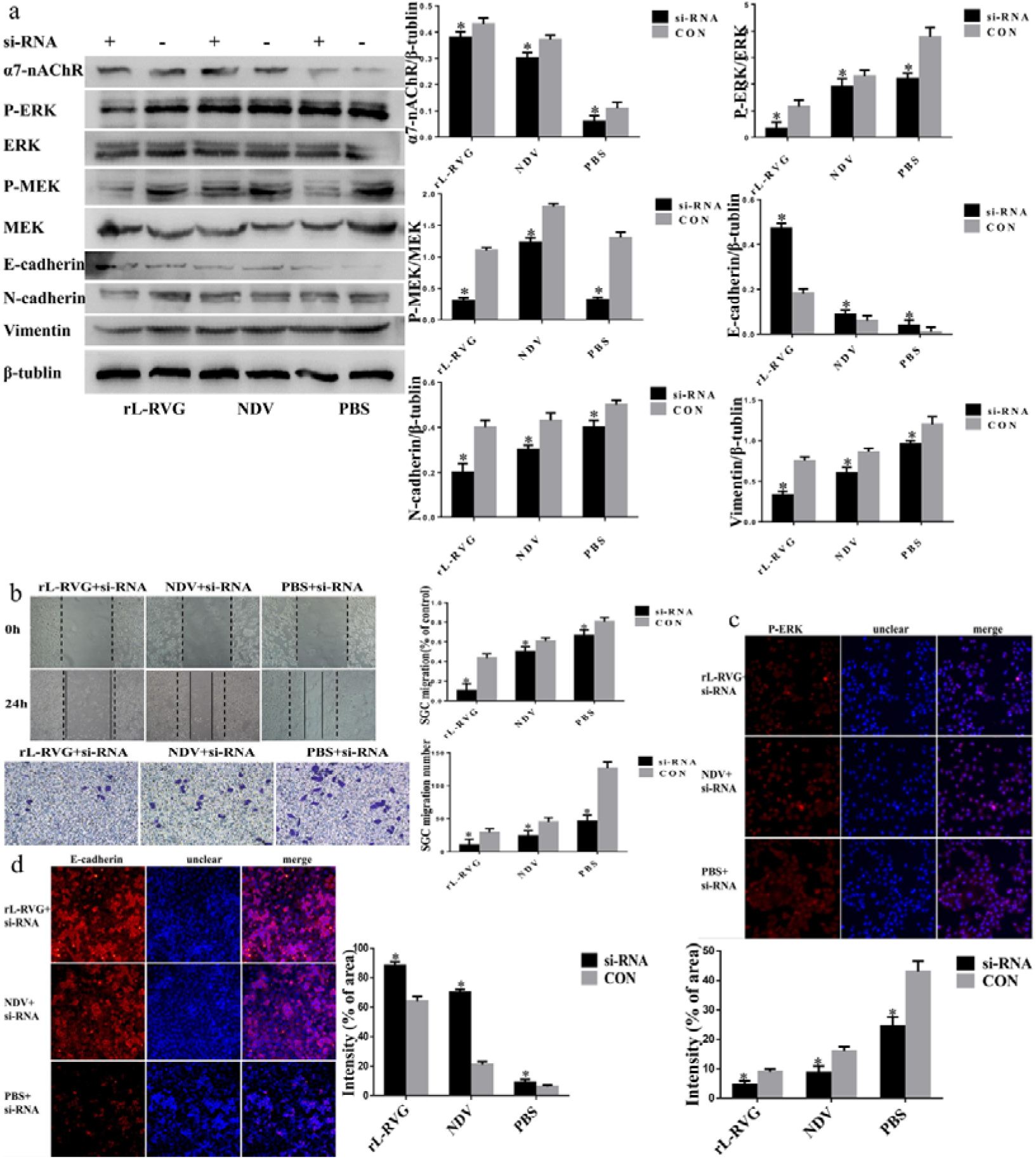
Effects of rL-RVG and si-RNA pretreated SGC cells on the α7-nAChR, MEK/ERK signaling pathway and epithelial/mesenchymal proteins and cell migration. **a**. Western blot analysis of α7-nAChR, MEK/ERK signaling pathway and epithelial/mesenchymal proteins. **b**. Cell migration was detected by wound healing and transwell assay. **c**. Immunofluorescence analysis of P-ERK. **d**. Immunofluorescence analysis of EMT proteins E-cadherin in infected SGC cells.*\*P<0.5, **P<0.01* (rL-RVG+si-RNA vs rL-RVG, NDV+si-RNA vs NDV, PBS+si-RNA vs PBS, Bar=25 μm).

**Figure 6.**
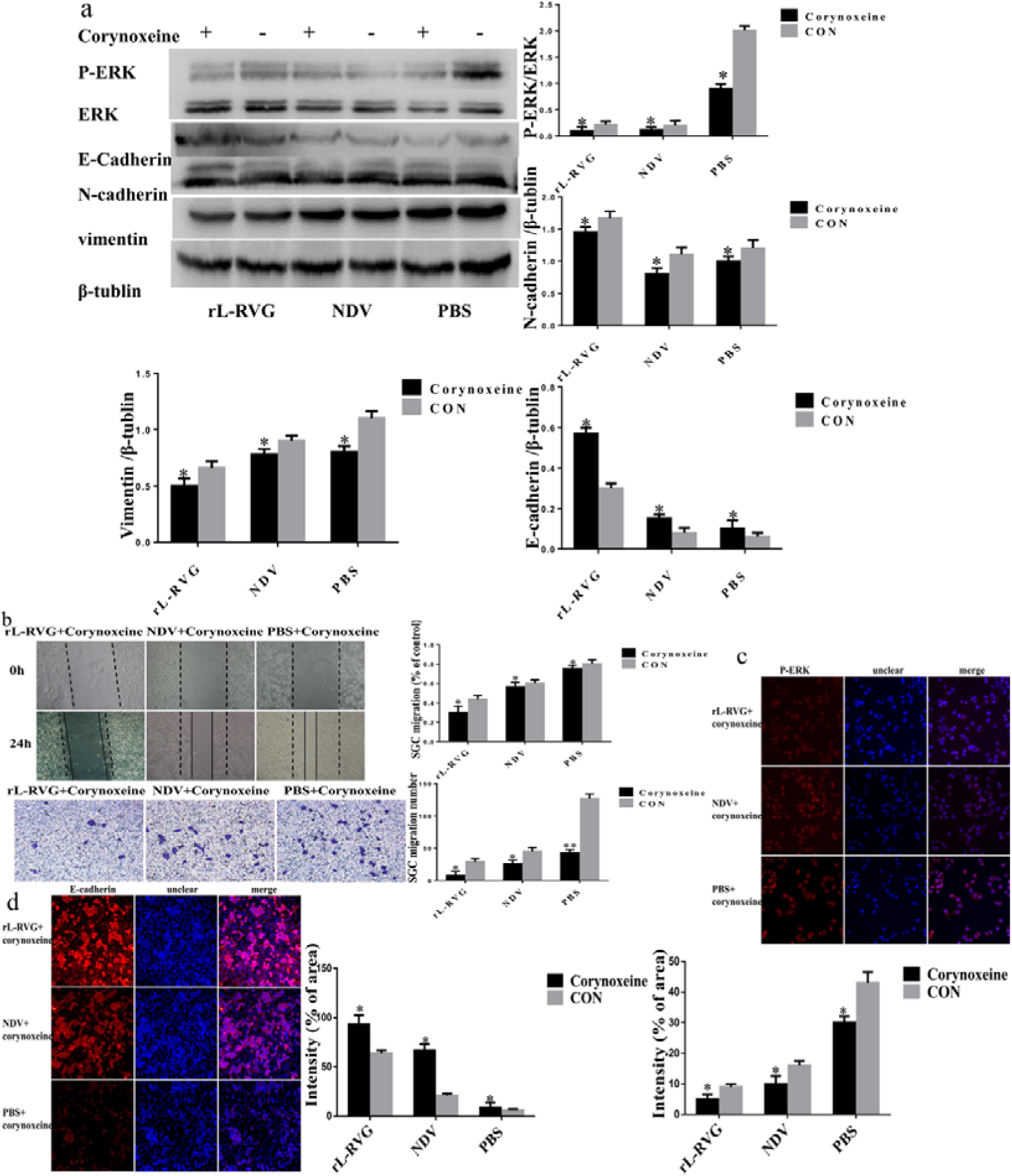
Effects of rL-RVG and corynoxeine pretreated SGC cells on α7-nAChR, ERK and epithelial/mesenchymal proteins and cell migration. **a**. Western blot analysis of α7-nAChR, P-ERK/ERK and epithelial/mesenchymal markers. **b**. Cell migration of SGC cells was detected by wound healing and transwell assay. **c**. Immunofluorescence analysis of P-ERK in SGC cells. **d**. Immunofluorescence analysis of EMT protein E-cadherin in infected SGC cells.*\*P<0.5, **P<0.01* (rL-RVG+corynoxeine vs rL-RVG, NDV+corynoxeine vs NDV, PBS+corynoxeine vs PBS, Bar=25 μm).

### rL-RVG reduced EMT by regulating α7-nAChR

In our study Western blot analysis revealed that the protein expression level of E-cadherin was increased while N-cadherin, and Vimentin were decreased in the rL-RVG or NDV groups compared with the PBS treated control group (Fig 1A). Furthermore, the groups pre-treated with MLA showed that the expression of E-cadherin was lower and the expression of N-cadherin and Vimentin was higher compared with the non-pretreated groups using western blot ananlysis (Fig 4A,D). We found similar results after the pretreatment with the si-RNA of α7-nAChR (Fig 5A,D), as well as in the corynoxenine treated group (Fig 6A,D). In addition the opposite result also obtained in ACB pretreating groups compared with the groups pretreated with MLA or si-RNA.

### rL-RVG inhibits subcutaneous growth of gastric cancer cells

A Tumor-bearing mouse model was established to confirm the antitumor effect of rL-RVG *in vivo*. Our results show that the tumor size of nude mice infected with rL-RVG and NDV was smaller than in the PBS group, and within the treated groups the tumor size in the rL-RVG group was smaller than in the NDV group (Fig. 7).

**Figure 7.**
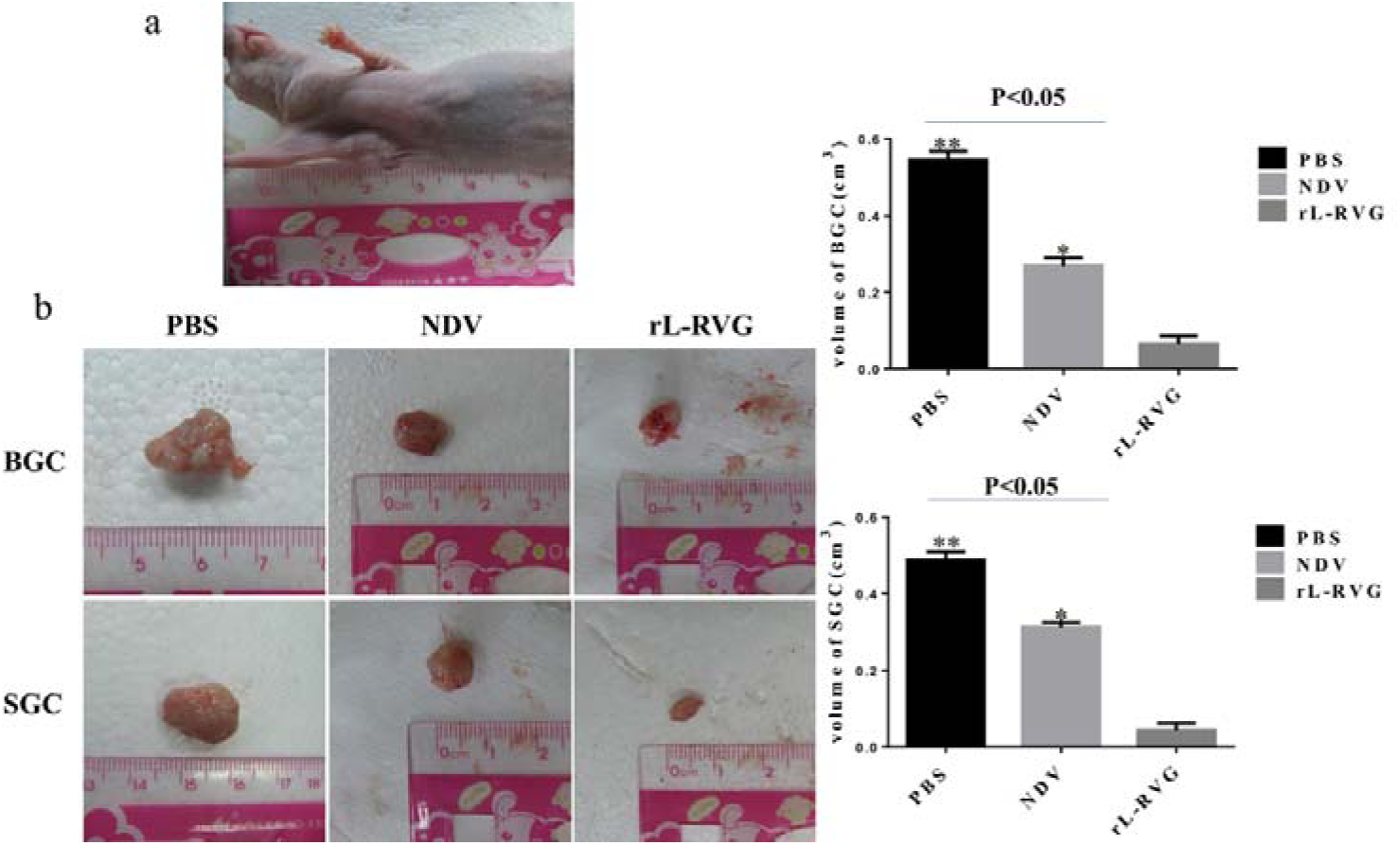
rL-RVG suppressed the growth of subcutaneous tumor of BGC and SGC cells in vivo. **a**. The subcutaneous tumor model was successfully established. **b**. BGC and SGC cells were pretreated with rL-RVG, NDV or PBS for 24 h and then subcutaneously injected into axillary subcutaneous tissues of adult female athymic-nude mice. The subcutaneous tumor was examined after two weeks post treatment. *\*P<0.5*, *\*\*P<0.01*.(rL-RVG vs NDV and PBS groups, respectively).

## Discussion

Although the incidence of gastric cancer has declined rapidly ^[17]^, it is still a major challenge to obtain early diagnosis and provide effective treatment options^[18]^. Tumor metastasis is the main cause of death in patients with gastric cancer. Therefore it is important to understand how to reduce metastasis formation and to find key methods of curing advanced-stage gastric cancer. In our study, recombinant avirulent NDV LaSota strain expressing the rabies virus glycoprotein rL-RVG did not only suppress migration of gastric cancer cells such as BGC and SGC cells *in vitro*, but also inhibited growth of subcutaneous tumor in nude mice *in vivo*. Our results demonstrate that rL-RVG suppresses gastric cancer cell migration by inhibiting the α7-nAChR/MEK-ERK-EMT axis.

With the development of modern cancer treatment, oncolytic therapy is becoming a novel biological treatment option that integrates gene therapy with immunotherapy. Recent studies revealed that the oncolytic Newcastle disease virus (NDV) replicates selectively and destroys tumor cells without damaging healthy cells^[19,20]^. Thus, selective killing has a higher efficiency accompanied by lower side effects. The 198-214 amino acid sequence of Rabies virus glycoprotein (RVG) is highly homologous with the 30–56 amino acid sequence in λ-bungarotoxin, which binds nAChRs. Therefore RVG has similar inhibiting effects as λ-bungarotoxin and has not only oncolytic effects but also blocks nAChRs by competitive antagonism which is a common mechanism of many pharmacological drugs. Thus, rL-RVG plays an role as competitive antagonist of α7-nAChR after infecting gastric cancer cell lines such as BGC and SGC.

nAChR consists of five subunits and assembles into heteromeric or homomeric pentamers. α7-nAChR, another type of nAChR, has a positive effect on cancer cell migration^[21]^. Nicotine and NNK could act as agonisst of α7-nAChR to facilitate the migration of gastric cancer cells^[12,22]^ and could indirectly activate ERK signaling through promoting the release of epidermal growth factor (EGF) and trans-activation of EGF receptors^[23]^. Moreover, the ERK signaling pathway also has influence on EMT that may regulate expression of mesenchymal and epithelial repressor genes^[24]^. In our study, we suggest that rL-RVG could lower the phosphorylation levels of ERK signaling and decrease EMT in SGC and BGC cells, indicating that gastric cancer cell migration which is suppressed by rL-RVG, is associated with the MEK-ERK signaling pathway.

It is of great significance to activate EMT for invasion and metastasis of gastric cancer cells^[25]^. An aberrant EMT activation typically results in the transformation of epithelial cells into mesenchymal cells and leads to phenotypic changes such as the loss of cell-cell adhesion, cell polarity and acquisition of migratory and invasive properties of cells. Cadherin is a significant component of adherent cell junctions. On one hand, aberrant activation of EMT could transform E-cadherin to N-cadherin, which is typically found in mesenchymal cells and could promote the formation of adhesions between cells and the stroma^[26]^. On the other hand, Vimentin is a widespread mesenchymal intermediate filament which results in adhesion and migration in activated cells ^[27]^ and is a hallmark of aberrant EMT activation in gastric cancer^[28]^. Our previous studies suggest that rL-RVG promotes apoptosis of gastric cancer cells^[14,29]^ but the effect of rL-RVG on gastric cancer migration and its underlying mechanism remain unknown. In this study, our results show for the first time that gastric cancer cell migration is suppressed after infection of gastric cancer cells with rL-RVG through competitive inhibition of α7-nAChR. This resulted in the decreased expression of mesenchymal markers including N-cadherin and vimentin and increased levels of the epithelial marker E-cadherin. Nicotine promotes migration of gastric cancer cells via the α7-nAChR pathway^[12]^. Thus we hypothesized that rL-RVG might suppress the migration of cancer cells via α7-nAChR. To demonstrate this we used ACB, MLA and si-RNA of α7-nAChR to pre-treat SGC cells and our results further confirmed our hypothesis.

Previous studies suggest that the ERK signaling pathway plays an important role in the proliferation, migration and apoptosis of cells^[30-31]^. Therefore, inhibiting the ERK signaling pathway strengthens the anti-tumor activity of gimatecan in gastric cancer^[32]^. Activation of ERK signaling results in the promotion of cervical cancer cell growth and metastasis^[33]^. Interestingly, we found that rL-RVG reduces the phosphorylation levels of MEK/ERK and rseulted in a decrease of phosphorylation levels in MLA, α7-nAChR and si-RNA pre-treated groups compared to the ACB pre-treated group. Additionally, we also achieved similar results in the change of EMT and migration of SGC cells. Hawsawi O and Henderson V suggested that HMGA2 may induce EMT via ERK signaling pathways^[34]^. In our study, after pretreating with corynoxenine, we found that the expression of N-cadherin and Vimentin was down-regulated and the expression of E-cadherin was upregulated compared to non-pretreated groups. We also found that rL-RVG has a much more inhibitory effect than NDV. Thus, rL-RVG does not only have the same effect as NDV, but also plays a role as competitive antagonist to inhibit α7-nAChR.

In conclusion, our research provides a new option in treating gastric cancer and provides insights into the mechanisms of rL-RVG on gastric cancer. For the first time we demonstrated that rL-RVG acts as competitive antagonist of α7-nAChR to suppress the migration of gastric cancer cells through inhibition of α7-nAChR-MEK/ERK-EMT.

